# Simultaneous detection of DNA and RNA virus species involved in bovine respiratory disease by PCR-free rapid tagmentation-based library preparation and MinION nanopore sequencing

**DOI:** 10.1101/269936

**Authors:** Matthew S. McCabe, Paul Cormican, Dayle Johnston, Bernadette Earley

## Abstract

The Oxford Nanopore MinION Mk1B is a portable 90 g device that sequences DNA directly at 450 bases/second generating sequence reads in excess of 400 kb. Recent improvements in error rate and speed of library preparation mean that this device has considerable potential for rapid molecular bovine pathogen diagnostics. We tested the MinION for rapid untargeted detection of viral pathogens associated with bovine respiratory disease (BRD), an economically important disease often involving primary infection of the lung by one or more of a number of DNA and/or RNA viruses. We combined three foetal lung cell cultures which were infected with either Bovine Respiratory Syncytial Virus (BRSV), Bovine Herpes Virus 1 (BoHV1) or Bovine Parainfluenza Virus 3 (BPI-3). BoHV1 is a DNA virus and BPI-3 and BRSV are RNA viruses. The cell cultures were treated with DNase and RNase to deplete bovine nucleic acid prior to viral nucleic acid extraction and double-stranded cDNA synthesis. Sequencing libraries were generated by PCR-free tagmentation in under 10 minutes and loaded onto a MinION sequencer. Approximately 7,000 sequencing reads were generated and analysed using high-throughput local BLAST against the NCBI nr/nt database. The top BLAST hit for 2,937 of these reads was identified as a virus. Of these, 2,926 (99.6%) were correctly identified either as BoHV1, BRSV or BPI-3.

## Introduction

Bovine respiratory disease (BRD) is the most costly disease of beef cattle in North America and causes significant losses in most other cattle producing regions including Ireland [1-3]. It is thought that the majority of BRD cases involve primary infection of the lower respiratory tract (LRT) with one or more viruses which predisposes the LRT to secondary infection with a single or multiple bacterial species [4,5].

As viruses are the most common primary pathogen in BRD, vaccines containing BRD associated modified live viruses (MLVs) are widely used in the cattle industry. The MLVs that are included in the most recent BRD vaccines (e.g. Rispoval^®^4 (Merck) and Bovi-Shield Gold ^®^5 (Zoetis)) are Bovine Respiratory Syncytial Virus (BRSV), Bovine Herpes Virus 1 (BoHV1), Bovine Parainfluenza Virus 3 (BPI3). These MLVs have been selected based on viruses that have been most commonly isolated and cultured or detected by qPCR from nasal swabs, lung washes and lung lesions from BRD cases in recent decades.

The primers and probes used in the qPCR assays are designed against small fragments (approximately 200 bp) of genomes of known (i.e. cultured and isolated) BRD-associated viral genomes and individual assays have to be performed for each BRD virus. In the majority of BRD cases, no virus is detected either by culturing or by qPCR. For example, in a recent report^3^ qPCR assays for BoHV1, BVDV, BPI3, BRSV and Bovine Corona Virus detected virus in only 8.7% (391 out of 4444) of nasal swab samples from live animals displaying BRD symptoms. The viruses used in vaccines (i.e. BoHV1, BVDV, BPI3, BRSV) were detected in only 5.4% (241 of the 4444) samples. In addition, despite the wide use of MLV BRD vaccines, BRD remains the leading cause of natural death in the cattle industry. This raises the possibility that other viruses or viral strains that are not easily cultured are involved in BRD and consequently targeted qPCR assays are an inadequate diagnostic method for BRD-associated viruses [6,7].

Recently viral metagenomics using next generation sequencing (NGS) platforms have been employed to survey viruses associated with BRD. This untargeted approach detects known and unknown viruses in a single universal assay. Ng *et al.* (2015) conducted viral metagenomic analysis of nasopharyngeal and pharyngeal recess swabs from animals with severe BRD symptoms and detected Bovine Adenovirus 3, Bovine Adeno-associated Virus, Bovine Rhinitis A Virus BSRI4, Bovine Rhinitis B Virus, Bovine Influenza D Virus, Bovine Astrovirus, Picobirnavirus, Bovine Parvovirus 2, and Bovine Herpesvirus 6. Bovine Adenovirus 3, Bovine Rhinitis A Virus BSR14 and Bovine Influenza D Virus BRSI1 were the only viruses that were significantly associated with BRD. None of the viruses that are included in current BRD vaccines were detected [7].

So far, BRD viral metagenomic studies have used Illumina sequencing by synthesis (SBS). This requires amplification of cDNA using techniques such as multiple displacement amplification (MDA), and library preparation that can take several hours. The read lengths on the Illumina MiSeq are short with a maximum of just 300 bp paired end on a 600 cycle reagent cartridge. SBS is relatively slow with a 600 cycle MiSeq run taking approximately 56 hours and no sequence data is available from a MiSeq until the run has completed. However, Illumina SBS platforms remain unsurpassed in terms of number and quality of sequence reads.

The MiSeq, which is one of the smaller Illumina NGS platforms (686 mm × 523 mm × 565 mm, 572 grams), is not designed for rapid diagnostics in the field. In contrast, the Oxford Nanopore Technologies MinION Mk1B is a pocket-sized (105 mm × 23 mm × 33 mm, 87 grams) field-deployable sequencing device that is based on nanopore sequencing. DNA is sequenced directly when it passes through recombinant *E. coli* CcsG nanopores that are embedded in a membrane and each base causes a characteristic change in the membrane current. This allows extremely rapid direct sequencing of individual DNA molecules. In October 2016 the R9.4 MinION flowcell was released which can run at 450 bases/second per nanopore and generate 10 gigabases of data per MinION flow cell. The library preparation with the ‘Rapid Sequencing Kit’ takes approximately 10 minutes. Loading the library takes approximately 20 minutes and thousands of long sequence reads (maximum length is typically >65 kb) are available for analysis within minutes of loading the library. Unlike the Illumina SBS platforms, the quality of nanopore reads does not decline with length. However, the error rate of reads from the MinION is still far higher than those of Illumina SBS platforms such as the MiSeq.

Several labs have used the MinION to detect viruses. Greninger *et al* (2015) reported untargeted metagenomic detection of high titre Chikungunya Virus (CHIKV), Ebola Virus (EBOV), and hepatitis C virus (HCV) from four human blood samples by MinION nanopore sequencing [8]. However, to obtain ≥1 μg of metagenomic complementary DNA (cDNA) for the library required for the nanopore sequencing protocol, randomly amplified cDNA was generated using a primer-extension pre-amplification method (Round A/B). Briefly, in Round A, RNA was reverse-transcribed with SuperScript III reverse transcriptase using Sol-PrimerA, followed by second strand synthesis with DNA polymerase with Sol-PrimerB and PCR amplification (25 cycles). Libraries were then prepared by end repair, adenylation and adapter ligation. After nanopore sequencing on the MinION they obtained sequences with an individual error rate of 24%. Despite the high error rate this allowed identification of the correct viral strain in all four isolates, and 90% of the genome of CHIKV was recovered with 97‒99% accuracy. Quick *et al.* (2016) identified high titre Ebola Virus in samples submitted less than 24 h after collection, with a targeted sequencing approach based on Ebola specific PCR primers that took 15-60 min [9]. For this they used targeted reverse transcriptase PCR (RT‒PCR) to isolate sufficient DNA for sequencing using a panel of 38 primer pairs that spanned the Ebola Virus genome. Recently, Kilianski *et al.* (2016) generated unamplified RNA/cDNA hybrids from nucleic acid extracted from either Venezuelan Equine Encephalitis Virus or Ebola Virus cell culture (both RNA viruses) and sequenced them individually on separate sequencing runs on a MinION. They were able to correctly identify each of the RNA viruses following alignment to the respective viral genomes [10].

In the present study, we tested the potential of untargeted nanopore sequencing on the MinION Mk1B for rapid simultaneous identification of a mixture of DNA and RNA viruses that are associated with BRD. Often more than one virus (with either a DNA or RNA genome) causes infection in the respiratory tract of an animal. We sequenced nucleic acid that we extracted from a mixture of three foetal lung cell cultures which were infected either with BRSV, BPI-3 or BoHV1 [11]. BoHV1 is a member of the family Alphaherpesviridae with a 150 kb linear double-stranded DNA monopartite genome. BRSV and BPI-3 are both members of the family Paramyxoviridae each with a 15 kb negative sense single-stranded RNA genome. BoHV1, BRSV and BPI-3 genomes are all packaged in a protein capsid which is surrounded by an outer lipid membrane envelope. BoHV1 also has a protein tegument between the capsid and the envelope.

We report correct simultaneous identification of combined DNA and RNA virus species involved in BRD by PCR-free rapid (10 min) tagmentation-based library preparation and nanopore sequencing on the portable Oxford Nanopore Technologies MinION Mk1B sequencer.

## Methods

### Viral cultures

Foetal lung cell cultures, infected with either BoHV1, BPI-3 or BRSV, were stored at −80°C. The cultures were sourced from (Agri-Food and Biosciences Institute (AFBI), Veterinary Science Division, Stormont, Belfast, N. Ireland). BPI-3 (TCID50 = 10^6.5^/100 μL in FCL) was cultured from a diagnostic lung sample. BHV-1 (TCID50 = 10^6.75^/100 μL in FCL) was cultured from a diagnostic lung sample. BRSV RISP_RS SP.C 11/03/15) (TCID50 = 10^3.75^/100 μL in FCL) is a vaccine strain of BRSV cultured in the FCL cell line.

### Nuclease treatment

The three frozen viral cultures were crushed to a fine powder with a sterile pestle and mortar under liquid nitrogen. Crushed frozen powder for each virus culture was weighed (BoHV1 (480 mg), BRSV (370 mg), BPI-3 (150 mg)) and combined in a 1.5 mL Eppendorf DNA low bind tube (Eppendorf, Hamburg, Germany). The volume was adjusted to 1 mL with DNA/RNA/DNase/RNase-free PBS (Sigma Aldrich Ltd., Arklow, Ireland) and 2.5 μL RNaseA (4 mg/mL) (Promega, Madison, WI, USA), 100 μL of 10× Turbo DNase buffer and 10 μL of Turbo DNase^TM^ (Life Technologies Ltd, Paisley, UK) were added. The solution was inverted slowly six times and incubated for 30 min at 37°C. Then another 10 μL of Turbo DNase was added, the mixture was inverted slowly again 6 times and incubated for a further 30 min.

### Nucleic acid extraction

Immediately following nuclease treatment, the remaining nucleic acids were extracted with the QIAamp Ultra Sens Virus Kit DNA extraction kit (Qiagen, UK) according to manufacturer’s instructions except that carrier RNA was substituted with 5.6 μL of a solution of 5 mg/mL linear acrylamide (Thermo Fisher Scientific, MA, USA). The final elution was performed with 2× 30 μL of buffer AVE (total 60 μL) which was supplied in the kit.

### Double-stranded cDNA synthesis

Double-stranded cDNA was generated with the Maxima H Minus Double-Stranded cDNA Synthesis Kit (Thermo Fisher Scientific, Waltham, MA, USA) according to the manufacturer’s instructions. Briefly, to generate the first cDNA strand (i.e. reverse transcription), 13 μL of extracted nucleic acid was added to 1 μL of random hexamer and mixed by gentle pipetting six times then incubated for 5 min at 65°C. The reaction was placed on ice for 1 minute, then centrifuged in a minifuge for 30 sec and placed back on ice. Then 5 μL of 4× First Strand Reaction Mix and 1 μL of First Strand Enzyme Mix were added, mixed by slowly pipetting the entire volume up and down six times, then incubated on an Eppendorf Master Cycler (Eppendorf, Hamburg, Germany) at 25°C (10 min), 50°C (30 min) and 85°C (5 min). The tube was removed from the Master Cycler and centrifuged for 10 seconds in a minifuge and placed on ice. To generate the second cDNA strand, the entire first strand cDNA reaction mixture (20 μL) was combined with molecular grade water (55 μL) (Sigma-Aldrich, Arklow, Ireland), 5× second strand reaction mix (20 μL) and second strand enzyme mix (5 μL). The entire volume (100 μL) was mixed by slowly pipetting up and down six times and was then incubated at 16°C (60 min). The reaction was stopped by adding 6 μL of 0.5 M EDTA (pH 8.0) (Sigma-Aldrich, Arklow, Ireland) and pipetting the entire volume up and down six times. To remove residual RNA, 10 μL (100 U) of RNase1 (supplied in the Maxima H Minus Double-Stranded cDNA Synthesis Kit) was added to the second strand reaction mixture which was mixed by slowly pipetting up and down six times and incubated at room temperature (22-25°C) for 5 min. The double-stranded cDNA reaction was purified using a Qiagen MinElute PCR clean up kit (Qiagen, Manchester, UK). This removed enzymes and primers and retained purified double-stranded cDNA and gDNA.

### Double-stranded cDNA/gDNA library preparation

The purified double-stranded cDNA/gDNA was tagmented using a Rapid Sequencing Kit SQK-RAD001 (Oxford Nanopore Technologies, Oxford, UK). Briefly, 7.5 μL of double-stranded cDNA/DNA was added to 7.5 μL of FRM (Oxford Nanopore Technologies, Oxford, UK), then mixed by slowly pipetting up and down six times. The reaction was incubated in a thermal cycler at 30°C for 1 min then at 75°C for 1 min. 1 μL of RAD (Oxford Nanopore Technologies, Oxford, UK) was added to the 15 μL tagmentation reaction and mixed by slow pipetting, then 0.2 μL of blunt/TA ligase master mix (New England BioLabs, Ipswich, MA, USA) was added to each tube and mixed again by slow pipetting. The reaction (designated pre-sequencing mix) was then incubated for 5 min at room temperature.

### Running the library

RAD (37.5 μL), H2O (31.5 μL) and pre-sequencing mix (6 μL) were combined in an Eppendorf LoBind tube (Eppendorf, Hamburg, Germany) and mixed by pipetting. The resulting 75 μL was loaded onto a Spot-on flowcell (FLO-MIN106) (Oxford Nanopore Technologies, Oxford, UK) on a MinION Mk1B (Oxford Nanopore Technologies, Oxford, UK) according to the manufacturer’s instructions. The flowcell was run for 16 h on MinKNOW software (Oxford Nanopore Technologies, Oxford, UK) using the Protocol ‘NC_48hr_Sequencing run_FLO-MIN105_plus_1D_Basecaller.py’.

### High throughput local BLAST search

The MinION generated 17,138 FAST5 sequence files which we converted to FASTA files using pore tools [12]. This resulted in 7,057 FASTA files that contained sequence reads. These sequence reads were then subjected to a local high throughput BLAST search against
the NCBI nr/nt database using a 24 core processor. The top hit BLAST result for each read was used for identification of the 7,057 sequences.

## Results

### Viral identification

Results of the high throughput BLAST search of the 7,057 MinION sequence reads are summarised in Table 1 and full details are included in S1 Table. A large number of these sequence reads (41.6%) were identified as viruses. The vast majority of the virus-identified sequence reads (99.6%) were identified as one of the three expected viruses BoHV1, BPI-3 and BRSV (Table 1). Only 11.6% of the sequence reads were identified as bovine and 46% of the sequence reads were identified as non-bovine/non-viral sequences.

**Table 1.** Summary of all top virus hits (species level) following high throughput local BLAST search of MinION sequence reads of rapid sequence library prepared from combined BPI-3, BRSV and BoHV1 calf foetal lung cell cultures against NCBI nr/nt database.

| Organism identified | Number<br>of reads | % of total<br>reads | % of virus reads |
| --- | --- | --- | --- |
| **BPI-3** (RNA virus, 15 kb genome) | 746 | 10.57 | 25.40 |
| **BRSV**(RNA virus, 15 kb genome) | 139 | 1.97 | 4.73 |
| **BoHV1** (DNA virus, 150 kb genome) | 2,041 | 28.93 | 69.5 |
| BoHV (no type designated) | 4 | 0.06 | 0.14 |
| BoHV5 | 4 | 0.06 | 0.14 |
| Herpesvirus type1 | 1 | 0.01 | 0.03 |
| *Bos taurus* BCL2/adenovirus | 1 | 0.01 | 0.03 |
| Nile crocodilepox virus | 1 | 0.01 | 0.03 |
| Total number of sequence reads | 7,057 | - | - |
| Total number of virus sequence reads | 2,937 | 42.2 | - |

### Read lengths and alignments

Twenty five of the reads were >10 kb and the longest read was 93,542 bp. However, these very long reads only had very short alignments to sequences on the NCBI nr/nt data base (Fig 1). The read length of sequences for which the top BLAST hit was virus, were longer for BoHV1 than for BRSV and BPI-3 (Fig 1). BoHV1 also had the longest alignments out of all viral and non-viral reads (Fig 1). There were three very long viral reads (36,880 bp, 46,607 bp, and 63,234 bp) for which the top hit was either the BoHV1 complete genome or the BoHV1.2 complete genome. However, the alignment length of these long reads to these genomes was short (499 bp, 1, 739 bp, 1, 046 bp respectively) (Fig 1).

**Fig 1.**
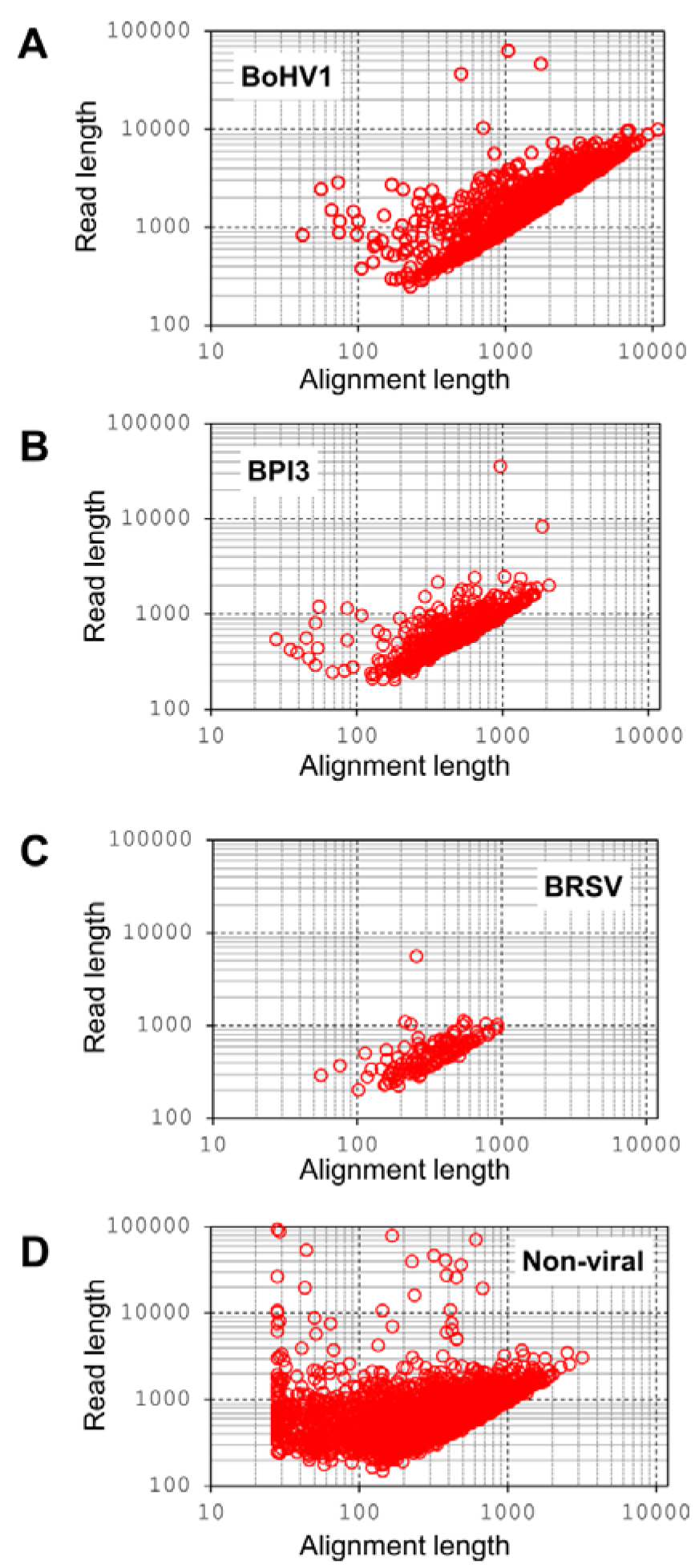
Scatterplot of alignment length (following high-throughput local BLAST search against NCBI nr/nt database) against read length of MinION sequence reads. Sequence reads were generated from a rapid sequencing library prepared from combined BPI-3, BRSV and BoHV1 calf foetal lung cell cultures. A: Reads for which the top BLAST hit was BoHV1. B: Reads for which the top BLAST hit was BPI-3. C: Reads for which the top BLAST hit was BRSV. D: Reads for which the top BLAST hit was not a virus.

The average percentage identity for alignments was 83.5% for viruses and 84.2% for non-viruses (Fig 2). Reads with >90% identity were <2 kb for both viruses and non-viruses (Fig 2).

**Fig 2.**
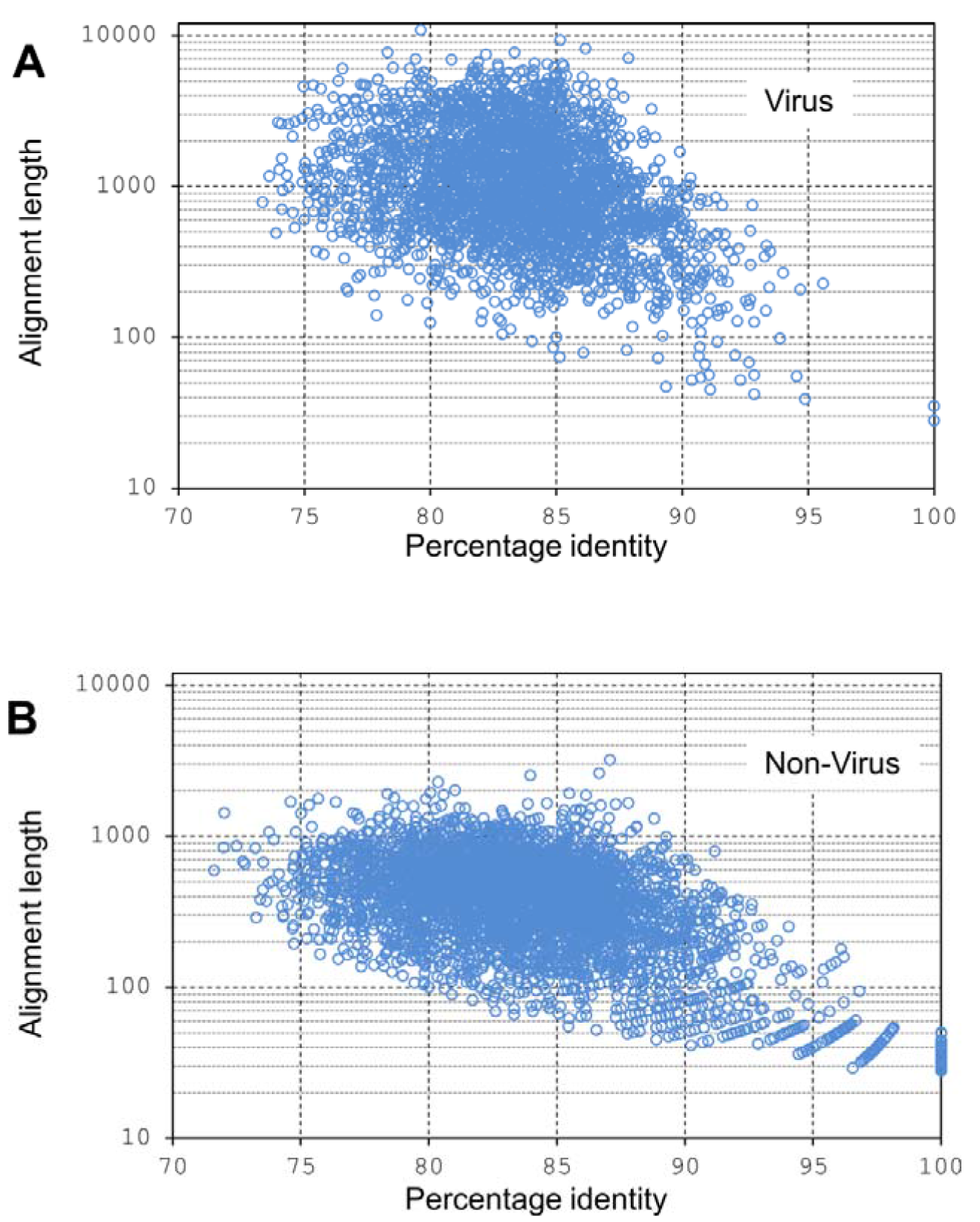
Scatterplot of alignment length against percentage identity of MinION sequence reads following high-throughput local BLAST against NCBI nr/nt database. Sequence reads were generated from a rapid sequencing library prepared from combined BPI-3, BRSV and BoHV1 calf foetal lung cell cultures A: Reads for which the top BLAST hit was virus B: Reads for which the top BLAST hit was not virus.

## Discussion

In the present study, combined DNA and RNA viruses were correctly identified following double-stranded cDNA synthesis, library preparation using the Oxford Nanopore Technologies Rapid Sequencing Kit, nanopore sequencing on a MinION Mk1B, and local high throughput BLAST against the entire NCBI nr/nt database. The rapid sequencing library preparation took approximately 10 min and, although the sequencer was run for 16 h, more than 5000 FAST5 files were generated in the first hour of sequencing, which was adequate for identification of the three viruses. This highlights the potential of the MinION for rapid diagnosis of viruses in regional veterinary laboratories and veterinary practices. As the MinION is highly portable, it also has potential for pen-side use by veterinarians and veterinary technicians to characterise viruses in animals that display BRD symptoms and this viral detection method could be applied to DNA extracted from lung washes or nasopharyngeal swabs. However, the numbers of reads generated for the nuclease-treated viral cell cultures was only 7,057 which would be insufficient to detect low titre viruses directly in clinical matrices. With the current depth of sequence available on the MinION, the only way to detect low titre viruses with this device will be through further improvement in depletion of host DNA and unbiased amplification of the library.

For nucleic acid extraction and cDNA synthesis we used a spin column method with a microcentrifuge and thermocycler. Magnetic bead-based nucleic acid extraction would eliminate the need for a centrifuge in the field but bead-based extractions tend to be inefficient compared to spin columns. At present, to treat the sample with nucleases, extract the viral nucleic acid then prepare and purify double-stranded cDNA takes approximately 5 hours but there is scope to optimise and simplify these steps to significantly reduce this time. We used our own high-throughput BLAST pipeline against the entire NCBI nr/nt data base for 7,057 reads. CPU time for this on a 24 core processor was 130 minutes. We also conducted a local BLAST search against only viral sequences (either against 90,000 virus sequences from NCBI or 7,000 sequences downloaded from Virusite) which took approximately 20 seconds. For virus-only BLAST searches all top hits were viral but many of these hits were not one of the three expected viruses. Reads that did not match the correct viruses were generally less than 100 bp, so removing short reads may work for virus-only BLAST. However, BLAST search against the entire NCBI nr/nt data base, though slower, appeared to be more robust as 99.6% of the viral top hits were correct to viral species level.

Due to the low ratio of viral nucleic acid to host nucleic acid, other viral metagenomics protocols commonly use whole genome amplification or PCR. However, in our study amplification was not necessary as 41.6% of the MinION sequence reads were identified as viral by BLAST search against the NCBI nr/nt data base. This may be due to the fact that we used stringent nuclease treatment with Turbo DNase which has 50× the activity of wild type DNase and that the titre of the virus was high in the cell cultures compared to clinical samples such as swabs. Other protocols also use spin filters to deplete eliminate host cells but it is not clear from the literature whether this is a benefit or an added complication that might bias the viruses that are detected [13]. Therefore we did not include a filtering step.

Interestingly the read lengths and alignment lengths were longer for the DNA virus (BoHV1) than the two RNA viruses (BPI-3 and BRSV). This is possibly due to the efficiency of the reverse transcriptase and the DNA polymerase in the double-stranded cDNA synthesis. As such, there may be bias towards DNA viruses in this protocol which will have to be allowed for if accurate quantification of the viruses is required. Direct RNA sequencing (without prior cDNA synthesis) using nanopore technology was recently announced [14].
However, for truly universal virus detection by sequencing we require direct DNA and RNA sequencing on the same flowcell which is not currently available.

The average error rate for 1D rapid sequencing libraries is currently 15% which explains the low percentage alignments to viral sequences on the NCBI nr/nt database that we observed. Oxford Nanopore Technologies claim this error rate will decrease to 5% for the rapid 1D library preparation and will be 1% for the slower 2D library preps by the middle of 2017. Surprisingly, despite the current high 15% error rate and low alignment percentages, our study showed that 99.6% of the MinION sequencing reads, for which the top BLAST hit was a virus, were correctly identified to species level. Several different viral strains were detected for BoHV1. (e.g. BoHV1 strain Cooper, BoHV1 subtype1 and BoHV subtype2). BoHV1 was a field isolate so it is possible that several strains were present. However, with its current error rate it is unlikely that MinION Mk1B 1D rapid sequencing can distinguish between viral strains or subtypes but as error rates are constantly improving this will likely be resolved in the near future.

We conclude that double-stranded cDNA synthesis, PCR-free tagmentation, nanopore sequencing on a MinION Mk1B with a 9.4 flowcell, and high-throughput local BLAST search can be used for rapid simultaneous species level identification of mixed RNA and DNA BRD-associated viruses in high titre cell cultures.

## Acknowledgments

We gratefully acknowledge Catherine Duffy and Michael McMenamy (Agri-Food and Biosciences Institute (AFBI), Stormont, Northern Ireland) for provision of viral cultures.

## Supporting information

**S1 Table. Results of local BLAST search of MinION Mk1B sequence reads against the NCBI nr/nt database**.

